# Interpretation of biological experiments changes with evolution of Gene Ontology and its annotations

**DOI:** 10.1101/228080

**Authors:** Aurelie Tomczak, Jonathan M. Mortensen, Rainer Winnenburg, Charles Liu, Dominique T. Alessi, Varsha Swamy, Francesco Vallania, Shane Lofgren, Winston Haynes, Nigam H. Shah, Mark A. Musen, Purvesh Khatri

## Abstract

Gene Ontology (GO) enrichment analysis is ubiquitously used for interpreting high throughput molecular data and generating hypotheses about underlying biological phenomena of experiments. However, the two building blocks of this analysis — the ontology and the annotations — evolve rapidly. We used gene signatures derived from 104 disease analyses to systematically evaluate how enrichment analysis results were affected by evolution of the GO over a decade. We found low consistency between enrichment analyses results obtained with early and more recent GO versions. Furthermore, there continues to be strong annotation bias in the GO annotations where 58% of the annotations are for 16% of the human genes. Our analysis suggests that GO evolution may have affected the interpretation and possibly reproducibility of experiments over time. Hence, researchers must exercise caution when interpreting GO enrichment analyses and should reexamine previous analyses with the most recent GO version.

## INTRODUCTION

Ontologies provide a uniform vocabulary for representing domain knowledge. The Gene Ontology (GO) is the most widely used ontology for specifying cellular location, molecular function, and biological process participation of human and model organism genes ^1^. The two building blocks of the GO are [1] the ontology itself and [2] the GO annotation. The ontology is a tree-like hierarchical structure of concepts (called GO terms) and their relationships to each other. The GO annotation is the list of all annotated genes linked to ontology terms describing those genes. The GO annotation documents all evidence that led to the association of a gene and a GO term by using evidence codes. In January 2015, 57% of all human gene annotations were assigned by automated methods, without curatorial judgment. They were labeled “inferred from electronic annotation” with the evidence code IEA. The remaining 43% of the gene annotations were manually assigned from experimental or computational analysis and author or curatorial statements. Manually assigned annotations are generally considered to be of better quality ^2^. Both the GO and its annotations are continuously evolving ^3–5^ as more experimental data become available. However, the human genome annotation is still incomplete. Furthermore, previous GO annotation versions were shown to be affected by confounds and biases such as annotation bias, where most annotations are for only few well-studied genes ^6,7^ or literature bias, where few articles disproportionally contribute many experimental annotations ^8^. These issues affect various applications relying on GO data including GO enrichment analysis ^5,9^, protein function prediction ^10,11^ or gene network analysis ^12,13^

The GO is predominantly used to analyze high-throughput data, such as gene expression microarray results. A typical analysis starts by identifying a list of differentially expressed genes. Then, to gain insight into the biological significance of the alterations in gene expression levels, researchers use GO enrichment analysis methods to determine whether GO terms about specific biological processes, molecular functions, or cellular components are over-or under-represented within the gene set of interest ^6^. Those methods can be based on different statistical methods and include traditional over-representation methods ^14,15^, Functional Class Scoring ^16,17^ or Pathway Topology ^18–20^. Wide adoption of the GO enrichment analysis in biomedical research is evident from tens of thousands of citations these tools have received. In many instances, enrichment analysis results are fundamental to the findings in research studies. For instance, Berry *et al.*^21^ concluded that there is “an interferon-inducible neutrophil-driven blood transcriptional signature in human tuberculosis,” where differentially expressed genes were identified to be interferon-related. The authors proposed a set of these genes as diagnostic markers. However, in subsequent years, it became clear that these genes, and interferon-stimulated genes in general, are not specific or sensitive to tuberculosis. Therefore, changes in the Gene Ontology and its annotations might affect the interpretation of experimental results ^5^.

There is an ongoing discussion about reproducibility of biomedical research that has significant effect on clinical translation ^22^. We hypothesized that use of continuously evolving ontologies may be one of the factors contributing to the reproducibility discussion because of evolving interpretation of the same data over time. To quantify the effects of GO evolution, we systematically analyzed 1.) the extent of changes in the Gene Ontology and the GO annotations between 2004 and 2015 and 2.) the effects of those changes on variation of p-values for enriched GO terms and on the consistency of GO enrichment analysis results over a decade. For this, we performed a Big Data analysis of gene sets derived from 104 multi-cohort analyses across 92 human diseases including more than 23,000 samples. Then, for each gene set, we systematically identified enriched GO terms ^23^ in specific versions of the ontology and annotations, monitored p-value changes over time, and quantified the consistency of a GO enrichment analysis result over time using the Jaccard Index ^24^. We found significant increases in the number of GO terms, annotated human genes, and annotations per gene. Furthermore, there was significant annotation bias in recent years as 58% of the annotations were for 16% of human genes. These significant changes in the GO and increased annotation bias resulted in very low consistency in enrichment analyses between earlier and later GO versions. Our analysis demonstrates that interpretations of the same gene set change over time with the evolution of the ontology and its annotations. This suggests that researchers must exercise caution when interpreting GO enrichment analyses.

## RESULTS

We analyzed two parameters. First, we looked at the extent of changes in the GO structure and in GO annotations between 2004 and 2015 by comparing the number of ontology terms and relationships, the number of GO annotations for the human genome, and the number of annotated human genes. Next, we analyzed the effects of those changes on the significance of p-values for enriched GO terms and on the overall consistency of GO enrichment analysis results (**Figure 1**).

**Figure 1.**
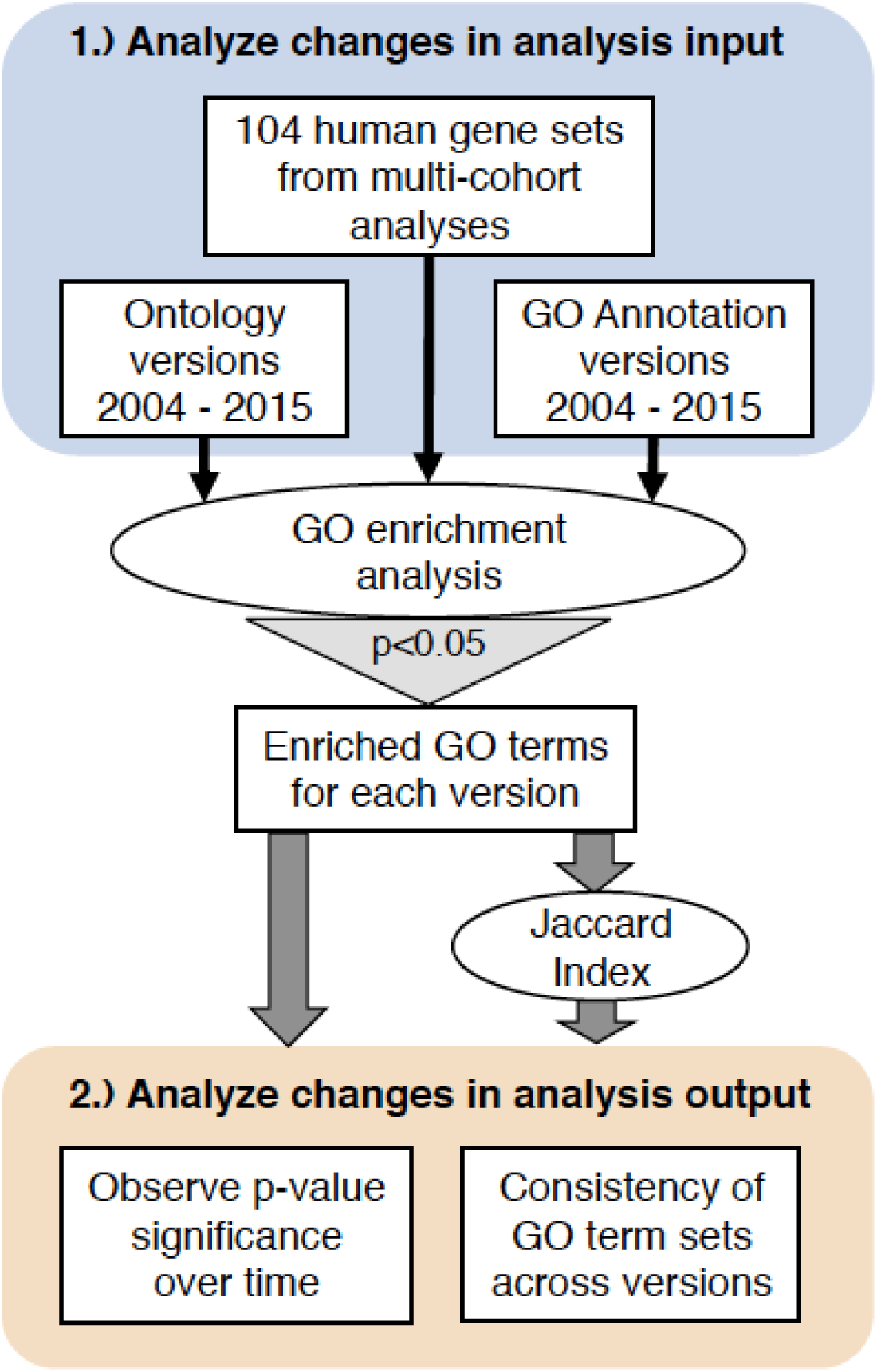
Overview of methods. We analyzed **1**) changes in input variables of GO enrichment analyses and **2**) how those changes affected enrichment analysis results over time.

### Quantifying the changes in GO during 2004 - 2015

We first analyzed the extent of changes between 2004 and 2015 in the Gene Ontology, GO annotations and human genes. We compared the number of ontology terms and relationships as well as the number of annotations describing the human genome. In the 11 years between February 2004 and January 2015, the number of terms in the GO increased by 2.5 folds (from 16,139 to 40,810; **Supplementary Figure 1A**), and the number of GO terms used for annotation of human genes increased by 3.8 folds (from 2,972 to 11,403; **Supplementary Figure and 1B**). In the same period, the number of relationships between terms increased by 3.5 folds (from 21,998 to 78,078; **Supplementary Figure 1A**). In the biological process and molecular function ontologies, which are the most informative ontologies for enrichment analyses, the increase was primarily due to 64,935 new relationships and 23,249 new GO terms. At the same time, 6,833 relationships were deleted, and 2,356 terms and 553 relationships were mapped to new terms and relationships (**Supplementary Figure 1C**). Further, the number of annotations increased by 6.3 folds (19,616 in 2004 to 109,162 in 2015; **Figure 2A**). Consequently, the proportion of protein-coding human genes with at least one GO annotation increased from 32% to 65%.

However, despite the increase in the number of annotated genes, the distribution of annotations remained skewed. We defined well-characterized genes as those with >10 GO annotations and poorly characterized genes as those with ≤10 annotations. Then, we compared the proportion of human genes with no, few (≤10) or many (>10) GO annotations. Only 16% of protein-coding human genes in 2015 had more than 10 annotations each (58% of the GO annotations), whereas 49% of protein-coding genes had 10 or fewer annotations (42% of GO annotations; **Figure 2A-B**). Importantly, one-third of protein-coding human genes still had no annotations. This bias persisted even when electronically inferred annotations were included in the analysis, where 69% of the annotations were for 27% of human genes (**Supplementary Figure 2**). It is possible that some of these genes lack annotations because curators have not yet reviewed the corresponding literature. However, the pervasive practice of forming new hypotheses based on enrichment analyses and selecting well-studied genes with many annotations for further study—and hence publication—is also responsible for this skew. For example, an 11-gene signature was shown to diagnose sepsis 2 to 5 days prior to clinical diagnosis ^25^, and it performed better than the current clinical tests ^26^. Unfortunately, 7 genes had <10 papers associated with them in NCBI Entrez Gene database. In contrast, >7,700 publications are associated with the tumor suppressor *TP53* in Entrez Gene. This discrepancy clearly shows the need for further experimental work to improve functional annotation of the human genome.

**Figure 2.**
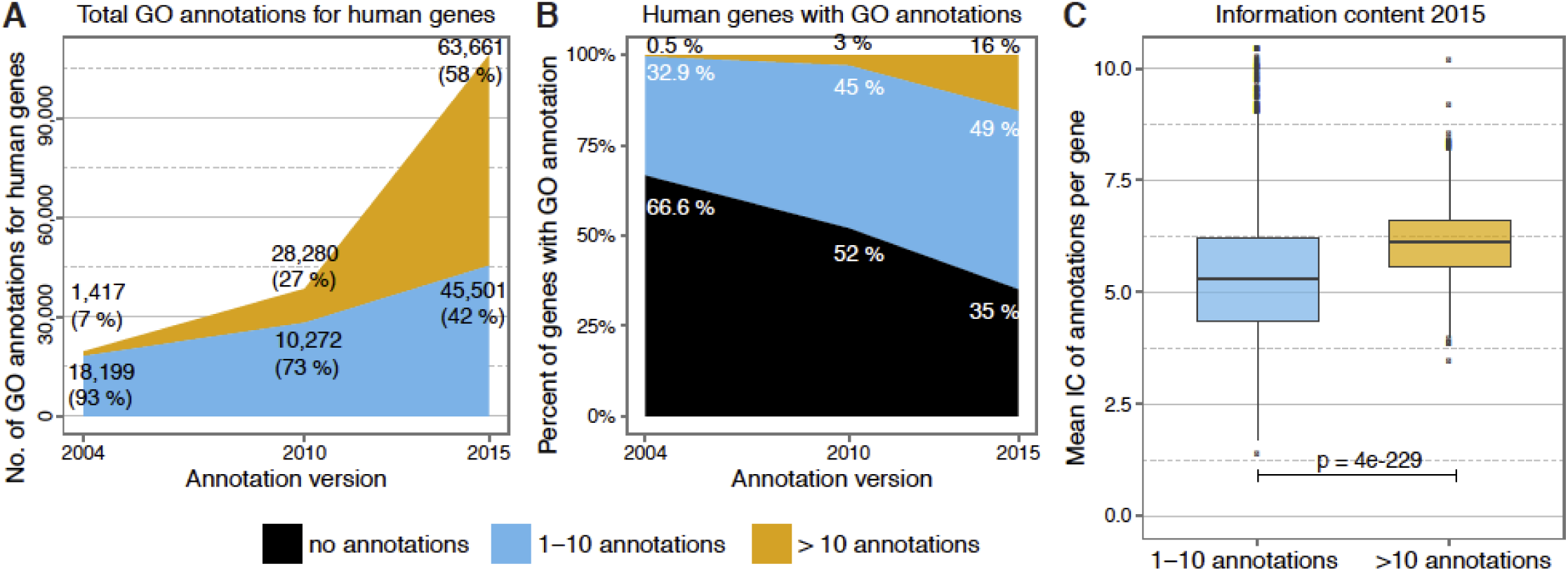
Gene ontology annotation developments, human genome, 2004 to 2015. **A**) Number of GO annotations and their distribution across poorly characterized (blue) and well-characterized (gold) human genes over time. **B**) GO annotation status of the human genome (2004 vs. 2015). Genes are classified by annotation status in uncharacterized (black) vs. poorly characterized (blue) vs. well characterized (gold). Only terms relevant for enrichment analysis results were counted (excluding: IEA, ND and cellular component). **C**) Comparison of the average information content (IC) of poorly characterized vs. well-characterized human genes in 2015 shows that the mean IC for genes with more annotations was higher (p = 4e-229). The same difference was observed in 2004 (p = 2e-19, Supplementary Figure 3).

We also measured bias by assessing the difference in mean information content (IC) of the annotations for less and more extensively studied genes. The IC of a GO term quantifies the specificity of the term in the context of the entire set of annotations. Terms annotating many genes are expected to be general and therefore have a low IC. Terms annotating only a few genes are specific, and have a high IC. Because the annotation count for a given term is up-propagated to all its parent terms, high-level terms always have a lower IC than their more specific child terms. We calculated the mean IC for less- and more extensively studied genes to quantify whether this annotation increase also constituted an increase in new information (i.e. precise and specific terms with high IC) or redundant information (e.g. adding only general terms with low IC). We found that the mean IC of extensively studied genes was higher than the mean IC of less-studied genes (p=4e-229; **Figure 2C**, **Supplementary Figure 3**), indicating that extensively studied genes have specific and detailed annotations, which were lacking in most of the less-studied genes. Collectively, these results illustrate that despite large increases in the Gene Ontology and GO annotations of human genes between 2004 and 2015, there is significant bias towards a small set of genes, which in turn can have significant impact on GO enrichment analysis, its consistency over time, and ultimately on interpretation of molecular data.

### Evolution of the GO affects the consistency of enrichment analysis results

To evaluate how changes in the GO and its annotations affect the interpretation of a list of differentially expressed genes, we collected whole genome expression profiles from more than 23,000 human samples across 377 independent datasets from 92 diseases. Then, for each disease, the we applied a multi-cohort analysis framework to identify disease gene signatures. This framework^27^ has been shown to identify reproducible gene signatures^28^ across multiple independent cohorts in different disease conditions, including bacterial and viral infections, organ transplantation, and cancer for identifying diagnostic and prognostic disease signatures, novel drug targets and repurposing FDA-approved drugs^25,28–33^. Next, we performed a GO enrichment analysis via traditional over-representation statistical methods for each disease gene signature, producing a set of enriched GO terms. We repeated this analysis using all combinations of historical Gene Ontology versions and GO annotation versions by year, and we monitored p-value changes over time. Furthermore, we calculated the consistency of GO enrichment analysis results for a given disease over time using the Jaccard Index^24^ as a measure of overlap between two sets of enriched GO terms to quantify changes in results as GO evolved. We chose Jaccard Index over other metrics such as the dice approach because of its robustness to low overlap between sets as well as to changes in set sizes, both of which are very likely to occur in our analysis. A consistency score of 0 indicates that two enrichment analysis results are entirely different, while a score of 1 indicates both analyses produced the same set of enriched terms. We used this score to explore changes in enrichment results across the complete set of disease gene signatures, using the March 2015 version of the annotation and ontology as our reference.

First, we compared each disease signature between different versions of the ontology while keeping the annotation version constant to January 2015 (the newest annotation independent of our reference). We observed an increase in median consistency from 0.27 in 2004 to 1 in 2015 (**Supplementary Figure 4A**). This large difference reflects significant re-structuring of the GO over the last decade. The steadiness of the trend also suggests the existence of small changes in the GO structure that were stable and propagated each year.

Next, we varied the annotation version while keeping the ontology version constant to January 2015 and observed low consistency until 2010 (median consistency range: 0.038 to 0.1), followed by a steady increase until 2015 (**Supplementary Figure 4B**). We observed the same trends when including electronic annotations (**Supplementary Figure 4C-D**) and across individual disease signatures (**Supplementary Figure 5A-D**). For instance, we analyzed gene signatures for influenza infection^33^ as well as pancreatic and non-small cell lung cancers, and observed trends in line with our general results. Collectively, these results suggest that changes in GO and its annotation have substantial effects on the results of enrichment analyses, which in turn can result in different interpretations for the same experiment depending on which version of the GO and its annotation were used for analysis.

### Evolution of the GO affects the p-value significance of enriched GO terms

Next, we analyzed the effects of annual GO and GO annotation updates on the significance of p-values for enriched GO terms to observe general trends for specific diseases. Thus, we changed both the ontology and the GO annotations together. For this test, we monitored p-value changes for all biological process terms that were deemed significant by at least one GO version for three diseases: influenza, non-small cell lung cancer, and pancreatic cancer (**Figure 3**). For the influenza gene signature, we hypothesized that child terms of the ontology branches *immune system process* or *response to stimulus* will be statistically significant as the stimulus induced by the influenza virus prompts an immune response in the host through transcriptional alteration of influenza signature genes. However, analysis of our influenza signature gene set returned at most 15 terms with p-values <0.05 before 2011, and none of the significant terms were child terms of those branches (**Figure 3A**). There are two reasons for lack of significance: (1) the term *immune system process* was added to the GO in September 2006, and (2) not enough genes were annotated with the terms in this branch. However, since 2012, many immune system- and stimulus-related terms became significant.

Similarly, for lung and pancreatic cancer gene signatures, we hypothesized that terms in the *cellular process* branch will be statistically significant because cell cycle-related events are essential to cancer survival (**Figure 3B-C**: purple). Unlike influenza, the GO enrichment analysis of the cancer signatures did not have an inflection point after which terms from the *cellular process* branch became significant. Instead, for the cancer gene signatures, we observed frequent changes between significant and insignificant p-values for individual GO terms, demonstrating that the enrichment results for cancer signatures were unstable over time. These results again demonstrate that different versions of the GO and its annotations could provide different interpretations of the same experiment.

**Figure 3.**
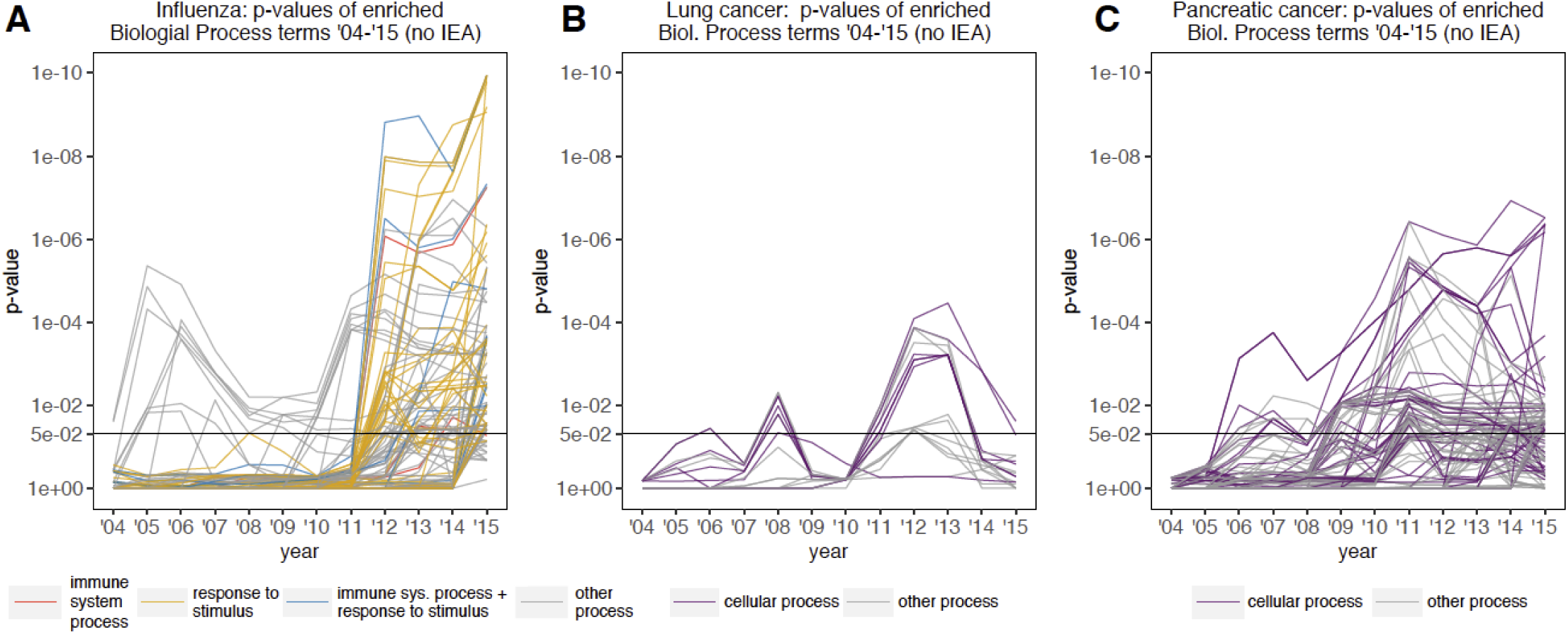
Significance of biological process GO terms over time with annual GO version updates (year of GO version = year of GO annotation version). Development of p-value significance in GO enrichment analysis result term sets in different GO versions are shown for subsets of significantly enriched biological process GO terms (p-value < 0.05 in at least one GO version) in three representative diseases: **A**) influenza, **B**) non-small cell lung cancer, and **C**) pancreatic cancer. Terms belonging to selected top-level branches in the biological process ontology are indicated in color (e.g. cellular process in violet).

### Evolution of GO affects the interpretation of enrichment analysis results

We explored how the interpretation of a disease signature would change with evolution of the GO, using the same three disease signatures as above. For each signature, we performed enrichment analyses using every possible pairing of ontology and annotation versions since 2004, and identified significantly enriched GO terms. In addition, we calculated the number of genes that were annotated with a term of interest, including any child terms, in the disease signature, and the reference gene set used for computing p-values over time. Finally, we calculated the IC of a GO term of interest to see whether bias—quantified via the IC—could affect interpretation of the analyses.

To test how well different GO versions represented current knowledge, we examined established disease mechanisms, which were expected to be identified in analyses of the selected diseases. First, we analyzed results from influenza infection. It has been known since 1981 that, upon infection, the host mounts a defense response by producing interferon-gamma ^34^ which, in turn, activates interferon-stimulated genes. We therefore expected the term *response to interferon-gamma* to be significant in our analysis. However, the term became enriched only when we used GO versions made starting in 2012 (**Figure 4A**). A number of factors contributed to this situation. First, the term *response to interferon-gamma* did not exist in the GO until March 2008. Second, once the term was introduced, it was annotated with very few genes (**Figure 4B**): up to 2011, only 15 genes were annotated with this term, of which only two were included in the 967 genes from our influenza infection signature. Consequentially, the IC for the term was high (>7.5) until 2011, indicating that it was used to annotate few genes (**Figure 4C**). As the number of genes annotated with *response to interferon-gamma* increased to 87 in 2012, its IC decreased to 5.6. This increase in annotations and genes for *response to interferon-gamma* is likely driven by research preference, as it coincided with increased research interest in influenza infection following the 2009 H1N1 influenza pandemic. A PubMed search revealed that in 2008, 2,824 influenza-related articles were published. This number increased to 5,586 in 2010. It is reasonable to assume that it took two years for this increased research output to be reflected in the GO annotations, when relevant terms correctly showed statistical significance for the gene signature. These observations underscore how before 2012 experimental influenza data may have been misinterpreted, or worse, deemed inconclusive and discarded.

**Figure 4.**
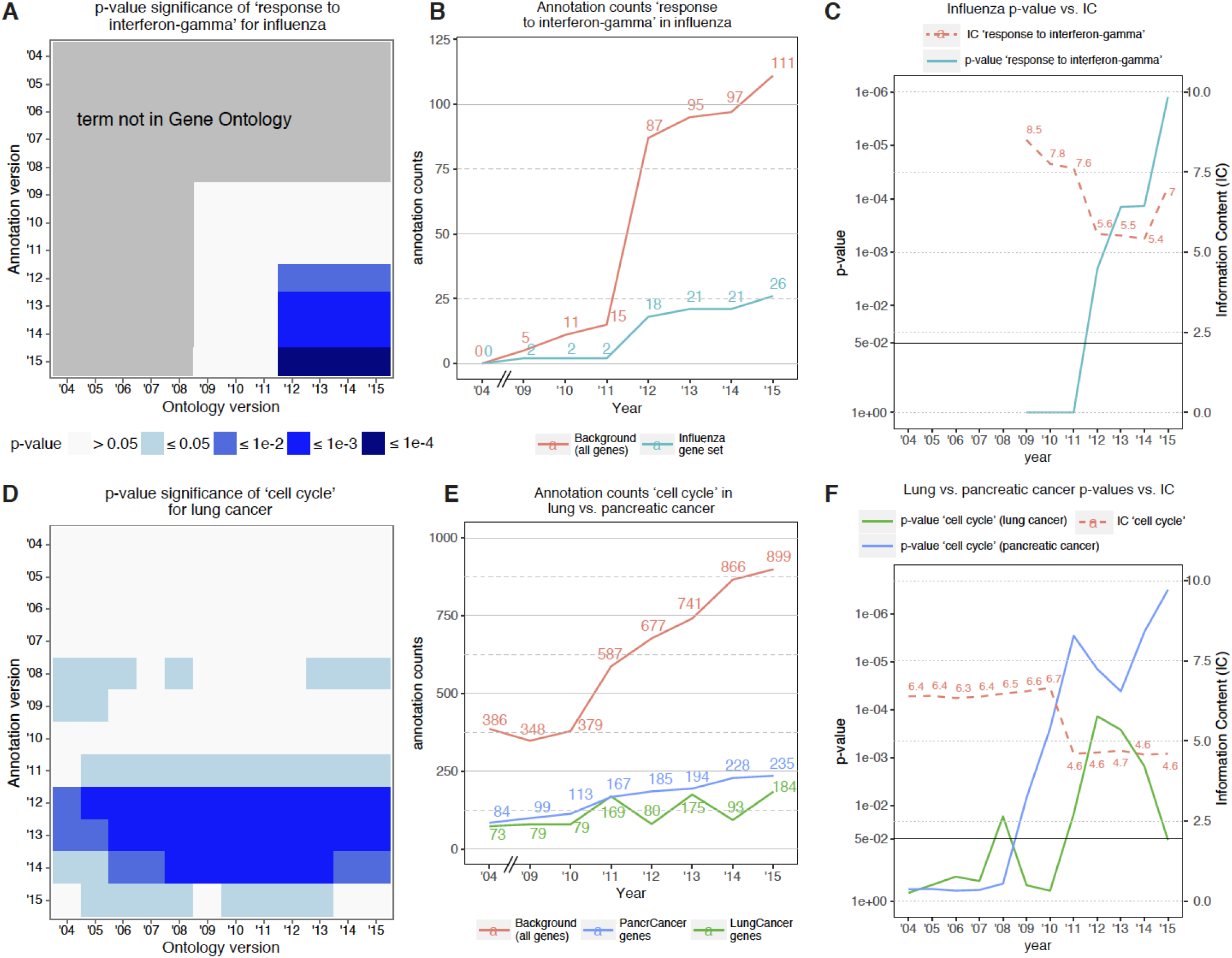
Effect of ontology and annotation version on consistency and significance of GO enrichment analysis results. **A**) Effect in influenza for GO terms: ‘response to interferon-gamma’. **B**) Number of human genes annotated with GO term: ‘response to interferon-gamma’ (including all child terms) in influenza gene set vs. background. **C**) Comparison of enrichment p-value and information content (IC) developments with annual updates of GO and GO annotations (year of GO version = year of GO annotation version) for ‘response to interferon-gamma’ in influenza**. D**) GO term enrichment significance for: cell cycle in non-small cell lung cancer (see Supplementary Figure 6 for pancreatic cancer) **E**) Number of human genes annotated with GO term cell cycle (including child terms) in pancreatic and non-small cell lung cancer gene sets vs. background (human genome). **F**) Comparison of enrichment p-value and IC developments with annual updates of GO and GO annotations for cell cycle in pancreatic and non-small cell lung cancer.

We observed similar results when we analyzed the term *cell cycle* in two cancer signatures. The role of dysregulated cell cycle in cancer has been well-established since the 1960s ^35^. The term *cell cycle* has existed in the GO since March 2001, and 386 human genes were annotated with the term (or any child terms) in 2004. However, *cell cycle* was not significant for either of the cancer gene signatures until 2008 (**Figure 4D, Supplementary Figure 6**), or until 2007 annotations if electronic annotations were included. However, in contrast to *response to interferon-gamma, cell cycle* is a more general term. Its IC from 2004 to 2010 was 6.5 (range: 6.3 to 6.7), which dropped to 4.6 in 2011 as the number of genes annotated with this term or any of its child terms increased from 379 to 587 in 2011, and has remained constant since then (**Figure 4E**). As more genes were annotated with *cell cycle*, we observed an increase in the significance of the term (**Figure 4E**). Interestingly, while the term continued to be statistically significant for the pancreatic cancer signature, it was not significant for the lung cancer signature **(Figure 4F**). This trend was not affected even when electronic annotations were considered. These results again illustrate how interpretation of a gene set could change with evolution of the GO and its annotations.

## DISCUSSION

GO enrichment analysis is virtually a *de facto* standard for interpreting high throughput molecular data and identifying underlying biological themes. GO evolution influences the results of enrichment analyses and interpretation of an experiment. Our analyses found increases in the number of GO terms, the number of annotated human genes, and the number of annotations for the human genome between 2004 and 2015. Gillis *et al.*^7^ previously investigated the stability of each gene’s ‘functional identity’ (agreement of gene-associated GO terms) over a 10-year period from 2001 to 2011, and they found that annotation bias in GO generally increased in humans. Our results are in agreement with those findings. Our analysis extend these results to show that the bias continues to increase rapidly also after 2011. In 2015, we found that 58% to 69% of the annotations are for 16% to 27% of the human genome, depending on whether electronically inferred annotations are included or not. For the same set of gene signatures, these changes and the bias in GO annotations may contribute to low consistency and different interpretations for enrichment results obtained using early and more recent GO versions.

Because of the lack of a gold standard for GO enrichment analysis, we used the latest (March 2015) version as reference to represent our current knowledge that is still incomplete and biased. To guard against this incompleteness and bias, we used established disease mechanisms that are *a priori* expected to be identified in analyses of the selected diseases. Yet, our analysis identified several factors that drive ontology evolution, which in turn could lead to misinterpretation, or worse no interpretation, of experimental data. For instance, a community effort to improve the representation of immunology content led to the introduction of 726 new GO terms covering immunological processes as well as revisions of existing immunology-related terms and relationships^36^. However, the number of human genes annotated with these terms remained low. For example, although the term *response to interferon-gamma* was added to GO in March 2008, only 15 genes were annotated with the term until 2011, which increased to 87 in 2012. The large increase is very likely due to increased efforts in influenza research following the influenza pandemic in 2009. This observation demonstrates that there is often a lag time until newly introduced terms are assigned to enough genes to become significant in enrichment analyses (4 years for *response to interferon-gamma).* Such a lag could in turn have further effect on interpretation of experiments.

Our results are in contrast with previous studies, which reported that despite evolution of GO, enrichment analyses are stable and do not necessarily change the interpretation ^5,9^. There are several important reasons for this discrepancy. First, the annotation bias was low in the GO annotations in 2010 compared to 2015 (**Figure 2**). Only 3% of the human genes had more than 10 annotations that accounted for 27% of annotations in 2010. Hence, prior to 2010, majority of the annotations (73%) were for less studied genes that restricted the effects of more studied genes on the enrichment analyses. In 2015, more studied genes represented 16% of the genes that accounted for 58% of annotations, and significantly enhanced the effect of annotation bias.

Second, Groß *et al.* used middle (2007) of the duration they analyzed (2003-2010) as the reference. The GO versions closer in time tend to have more significant categories in common (**Supplementary Figure 4**). Hence, choosing a reference version in the middle of the evaluation period is unlikely to observe cumulative effect of smaller changes aggregated over a longer period on interpretation of enrichment analyses. In contrast, we chose the last version of the GO (2015) as the reference for evaluating effect of GO evolution from 2004 to 2015, which allowed us to aggregate changes over more than a decade to observe effects of GO evolution on enrichment analyses.

Third, Groß *et al.* used the dice approach for computing stability, which corresponds to the harmonic mean and is suitable for comparing sets of similar size. In contrast, we used Jaccard index as a measure of stability, which produces slightly different results than dice, especially when overlap between two results is low^37^. The choice of a stability measure in turn is dependent of the amount of data used for analysis. Groß *et al.* only used two datasets and GO terms with at least 20 genes, which justified use of the dice approach. In contrast, we used 377 datasets composed of over 23,000 human samples to derive 104 gene signatures to perform systematic global analyses of effects of GO evolution. We also did not restrict the set of GO terms used in our analyses based on the number of genes, which further justified our use of Jaccard index. Arguably, a limitation of our analysis is that Jaccard index does not account for semantic similarities of ontology term. It is possible that accounting for semantic similarity may increase overall consistency for some diseases. However, different number of enriched terms between early and later versions of GO for a given gene signature demonstrate that the low consistency observed in our analysis is mostly independent of semantic similarity, and that the effect of accounting for semantic similarity will likely be limited. For instance, for the influenza gene signature, compared to 112 significant terms in 2015, there were only four significant terms in 2004, none of which were stimulus- or immune-related. Similarly, for pancreatic cancer, no terms were significant in 2004 compared to 81 significant terms in 2015. Further, given that 35% of the human genes still do not have any GO annotations (20% when including electronically inferred annotations), this problem is likely to persist for a foreseeable future. If the GO continues to grow at the same rate as in the last 5 years, it will likely still take more than 10 years before 90% of the human genes are annotated. In our analysis 21-23% of disease gene signatures (8-14% when including electronically inferred annotations) did not return any significant terms and therefore had to be excluded from further consistency analyses. Therefore, special attention must be paid when interpreting a gene list with a substantial number of un-annotated genes in the list.

In general, enrichment analysis results are intended as an exploratory approach to organize and probe large scattered datasets. Our analyses show, especially in early GO versions with a lot of missing annotation data, that enrichment results with many terms close to the significance cutoff (p-values ~ 0.05) can be noisy. In practice, this means that enrichment results should be filtered and curated by an expert in the topic under study before additional experiments are performed. Since we performed an analysis comparing a very large number of enrichment results with each other, we could not account for the expert curation step, which should be kept in mind when interpreting our results. Our study is an attempt to illustrate the extend and effects of GO evolution at a large scale and especially including recent GO versions, which were not included in earlier studies. Our results highlight the importance to develop methods for assessment of enrichment results if correctness is unknown due to still incomplete annotation data. Towards this goal, a recent study by Ballouz *et al.*^38^ demonstrated that robustness and uniqueness of enrichment results can be used as method for bias correction and for assessment whether enrichment analysis results are biologically meaningful.

Therefore, we suggest the following best practices for publication and re-use of GO enrichment analysis results. Publications should document the exact GO and annotation version used for analysis, and provide gene signature used for enrichment analysis in an easily accessible format to simplify re-running enrichment analyses once GO updates become available (e.g., CSV or text files using standard gene identifiers). Publications should also report the number and proportion of uncharacterized genes excluded from analysis, and include the number of well-studied genes in gene signature since they contribute most annotations. Further, it is advisable to periodically re-analyze a gene signature as recommended by the GO Consortium ^39^, and apply assessment methods to enrichment analysis results when correctness is unknown.

In summary, the ongoing discussion regarding reproducibility in biomedical research is justifiably focused on appropriate use of statistical methods and experimental models. However, our results strongly suggest that it should also include how using current knowledgebases, especially GO, to interpret experimental data is affected by evolution of these knowledgebases. It is very likely that changing interpretation of an experiment due to evolution of these knowledgebases could be viewed as irreproducible results. However, in these cases, it is important to highlight that it is the interpretation of data generated from an experiment that is not reproducible instead of the data itself; that the data themselves may be, and very likely are, still reproducible. Biomedical researchers must exercise caution when interpreting experimental results, and continue to reexamine previous analyses periodically with the most recent GO version.

## MATERIALS AND METHODS

### Gene ontology

We obtained archived versions of GO and GO annotations in annual intervals from 2004 to 2015 from the Gene Ontology website (ftp://ftp.geneontology.org/pub/go/ontology-archive, http://www.ebi.ac.uk/GOA/archive and and ftp://ftp.geneontology.org/pub/go/godatabase/archive/full/).

**Table 1.** **GO annotation and ontology versions used in this analysis.**

| Year (GO version) | '04 | '05 | '06 | '07 | '08 | '09 | '10 |
| --- | --- | --- | --- | --- | --- | --- | --- |
| Annotation version | 2004/03/29 | 2005/02/11 | 2006/01/19 | 2007/01/13 | 2008/01/20 | 2009/01/18 | 2010/01/24 |
| Ontology version | 2004/02/14 | 2005/01/03 | 2006/01/03 | 2007/01/03 | 2008/01/05 | 2009/01/09 | 2010/01/05 |
| Year (GO version) | '11 | '12 | '13 | '14 | '15 | 2015 Reference |  |
| Annotation version | 2011/01/01 | 2012/01/11 | 2013/01/08 | 2014/01/18 | 2015/01/08 | 2015/03/05 |  |
| Ontology version | 2011/01/05 | 2012/01/04 | 2013/01/05 | 2014/01/07 | 2015/01/07 | 2015/03/13 |  |

For all human genes, we generated GO annotation statistics by counting the number of enrichment-analysis-relevant GO annotations. We excluded terms with GO evidence codes Inferred from Electronic Annotation (IEA) or No biological Data available (ND), terms from cellular component ontology, and duplicate annotations that only differ in evidence codes. Changes between 2004 and 2015 Gene Ontology versions were calculated using the COntoDiff tool ^40^.

### Gene sets

Next, we obtained differentially expressed gene sets from the MetaSignature Database ^27^ (Version January 2015 http://metasignature.stanford.edu/), which contains 104 manually curated multi-cohort analyses for 92 diseases, with normalized expression levels derived from 377 individual microarray experiments (**Supplementary Dataset 1**). All datasets were downloaded from Gene Expression Omnibus (GEO). Multi-cohort analyses were performed using our Metaintegrator pipeline ^29^, and we applied a false discovery rate (FDR) cutoff of 0.01 to select sets of differentially expressed genes for each experiment.

GEO datasets used for influenza multi-cohort analysis ^33^: GSE6269, GSE52428, GSE42026, GSE40012, GSE389000, GSE38900, GSE34205, GSE32139, GSE32138, GSE20346, GSE17156

GEO datasets used for non-small cell lung cancer multi-cohort analysis: Bhattacharjee, GSE10072, GSE1037, GSE11969, GSE19188, GSE2514, GSE4824, GSE7670, Shedden

GEO datasets used for pancreatic cancer multi-cohort analysis: E-EMBL-6, E-MEXP-1121, E-MEXP-950, GSE11838, GSE15471, GSE15550, GSE16515, GSE19650, Sourtherland

### Gene symbol mapping

Mappings to gene symbols were not provided in GO annotation files before 2009. Therefore, we combined all GO annotation gene mappings from all years to account for the fact that earlier versions of GO annotations did not contain complete mappings. We removed annotations with obsolete identifiers used in early versions that could not be mapped to any current gene.

### Enrichment analysis

We analyzed the gene sets using BinGO, a Cytoscape plugin for GO enrichment analysis. BinGO applies traditional overrepresentation statistical methods to produce a set of enriched GO terms, including enrichment p-values.^14^ For each gene set, we re-ran the analysis using all possible combinations of GO and GO annotation versions, including and excluding electronic annotations. We recorded each GO term’s enrichment p-value across all ontology and annotation versions, producing a two-dimensional time series of the enrichment p-value for all GO terms. To create the result set of terms used to generate hypotheses, we selected terms with FDR corrected p-values <= 0.05 in the hypergeometric test.

### Consistency calculation

To quantify the consistency/overlap of the result set of an enrichment analysis over time, we used the Jaccard index^24^. A consistency score of 0 indicates that two sets of results are entirely different, while a score of 1 indicates that both enrichment analyses produced the same result set. We excluded consistency scores from gene sets that did not return any significant GO terms with the selected fixed 2015 version of annotation or ontology. The number of gene sets used for generation of consistency plots, is indicated in corresponding consistency plot headings.

### GO annotation counts

The term GO annotation count refers to the number of genes annotated with a certain term. This parameter was calculated by counting all genes annotated with a term of interest in a given GO annotation version, including all genes annotated with child terms of the term of interest.

### Calculating Information Content (IC) scores

We used the IC of a GO term as a proxy for its usage and specificity. The IC of a GO term ***t*** is defined as negative log likelihood IC(***t***) = −log_2_ ***P***(***t***), where ***P***(***t***) is the probability of finding ***t*** within all GO annotations for human genes of a given year. We calculated the IC for each GO term based on the number of human genes annotated with it (or any of its child terms in a given GO version) according to a given Gene Ontology Annotation (GOA) file. The IC quantifies the specificity of a term in the context of the entire set of annotations for human genes, where terms annotating many genes, such as *cell cycle*, are expected to be general terms and are assigned a low IC. Terms annotating only a few genes, such as *response to interferon-gamma,* are specific terms with a high IC. We calculated the IC for any term and any combination of Gene Ontology and Gene Ontology Annotation file for each year from 2004 to 2015 using the using ***Resnik IC*** implementation ^41^ of the Semantic Measures Library (SML) ^42^. Furthermore, we computed each gene’s mean IC as the mean IC of the GO terms assigned to it.

## Contributions

J.M.M., M.A.M. and P.J.K. conceived the project. A.T., J.M.M., D.T.A., C.L., F.V., W.H., S.L., V.S. and P.J.K. collected data and performed data analyses. A.T., J.M.M, R.W. and D.T.A. performed the experiments and analyzed data. A.T. and P.J.K. wrote the manuscript with input from all other authors.

## References

1. Ashburner, M. et al. Gene ontology: tool for the unification of biology. The Gene Ontology Consortium. Nat Genet 25, 25–9 (2000).

2. Schnoes, A. M., Brown, S. D., Dodevski, I. & Babbitt, P. C. Annotation error in public databases: misannotation of molecular function in enzyme superfamilies. PLoS Comput. Biol. 5, e1000605 (2009).

3. Huntley, R. P., Sawford, T., Martin, M. J. & O’Donovan, C. Understanding how and why the Gene Ontology and its annotations evolve: the GO within UniProt. GigaScience 3, (2014).

4. Bodenreider, O. & Stevens, R. Bio-ontologies: current trends and future directions. Brief. Bioinform. 7, 256–274 (2006).

5. Groß, A., Hartung, M., Prüfer, K., Kelso, J. & Rahm, E. Impact of ontology evolution on functional analyses. Bioinforma. Oxf. Engl. 28, 2671–2677 (2012).

6. Khatri, P., Sirota, M. & Butte, A. J. Ten years of pathway analysis: current approaches and outstanding challenges. PLoS Comput. Biol. 8, e1002375 (2012).

7. Gillis, J. & Pavlidis, P. Assessing identity, redundancy and confounds in Gene Ontology annotations over time. Bioinformatics 29, 476–482 (2013).

8. Schnoes, A. M., Ream, D. C., Thorman, A. W., Babbitt, P. C. & Friedberg, I. Biases in the Experimental Annotations of Protein Function and Their Effect on Our Understanding of Protein Function Space. PLoS Comput. Biol. 9, e1003063 (2013).

9. Clarke, E. L., Loguercio, S., Good, B. M. & Su, A. I. A task-based approach for Gene Ontology evaluation. J. Biomed. Semant. 4 Suppl 1, S4 (2013).

10. Jiang, Y., Clark, W. T., Friedberg, I. & Radivojac, P. The impact of incomplete knowledge on the evaluation of protein function prediction: a structured-output learning perspective. Bioinformatics 30, i609–i616 (2014).

11. Gillis, J. & Pavlidis, P. The impact of multifunctional genes on ‘guilt by association’ analysis. PloS One 6, e17258 (2011).

12. Gillis, J. & Pavlidis, P. ?Guilt by Association? Is the Exception Rather Than the Rule in Gene Networks. PLoS Comput. Biol. 8, e1002444 (2012).

13. Gillis, J., Ballouz, S. & Pavlidis, P. Bias tradeoffs in the creation and analysis of protein?protein interaction networks. J. Proteomics 100, 44–54 (2014).

14. Maere, S., Heymans, K. & Kuiper, M. BiNGO: a Cytoscape plugin to assess overrepresentation of gene ontology categories in biological networks. Bioinforma. Oxf. Engl. 21, 3448–3449 (2005).

15. Draghici, S. Onto-Tools, the toolkit of the modern biologist: Onto-Express, Onto-Compare, Onto-Design and Onto-Translate. Nucleic Acids Res. 31, 3775–3781 (2003).

16. Subramanian, A. et al. Gene set enrichment analysis: a knowledge-based approach for interpreting genome-wide expression profiles. Proc. Natl. Acad. Sci. U. S. A. 102, 15545–15550 (2005).

17. Dennis, G. et al. DAVID: Database for Annotation, Visualization, and Integrated Discovery. Genome Biol. 4, P3 (2003).

18. Draghici, S. et al. A systems biology approach for pathway level analysis. Genome Res. 17, 1537–1545 (2007).

19. Tarca, A. L. et al. A novel signaling pathway impact analysis. Bioinforma. Oxf. Engl. 25, 75–82 (2009).

20. Mi, H., Poudel, S., Muruganujan, A., Casagrande, J. T. & Thomas, P. D. PANTHER version 10: expanded protein families and functions, and analysis tools. Nucleic Acids Res. 44, D336–342 (2016).

21. Berry, M. P. R. et al. An interferon-inducible neutrophil-driven blood transcriptional signature in human tuberculosis. Nature 466, 973–977 (2010).

22. Begley, C. G. & Ellis, L. M. Drug development: Raise standards for preclinical cancer research. Nature 483, 531–533 (2012).

23. Khatri, P. & Drăghici, S. Ontological analysis of gene expression data: current tools, limitations, and open problems. Bioinforma. Oxf. Engl. 21, 3587–3595 (2005).

24. Jaccard, P. Étude comparative de la distribution florale dans une portion des Alpes et des Jura. Bull. Société Vaudoise Sci. Nat. 37, 547–579 (1901).

25. Sweeney, T. E., Shidham, A., Wong, H. R. & Khatri, P. A comprehensive time-course-based multicohort analysis of sepsis and sterile inflammation reveals a robust diagnostic gene set. Sci. Transl. Med. 7, 287ra71 (2015).

26. Sweeney, T. E., Wong, H. R. & Khatri, P. Robust classification of bacterial and viral infections via integrated host gene expression diagnostics. Sci. Transl. Med. 8, 346ra91 (2016).

27. Haynes, W. A. et al. ‘Empowering Multi-Cohort Gene Expression Analysis to Increase Reproducibility,’. PSB Sess. Methods Ensure Reprod. Biomed. Res. Rev.

28. Sweeney, T. E., Haynes, W. A., Vallania, F., Ioannidis, J. P. & Khatri, P. Methods to increase reproducibility in differential gene expression via meta-analysis. Nucleic Acids Res. gkw797 (2016). doi:10.1093/nar/gkw797

29. Khatri, P. et al. A common rejection module (CRM) for acute rejection across multiple organs identifies novel therapeutics for organ transplantation. J. Exp. Med. 210, 2205–2221 (2013).

30. Mazur, P. K. et al. SMYD3 links lysine methylation of MAP3K2 to Ras-driven cancer. Nature 510, 283–287 (2014).

31. Chen, R. et al. A meta-analysis of lung cancer gene expression identifies PTK7 as a survival gene in lung adenocarcinoma. Cancer Res. 74, 2892–2902 (2014).

32. Sweeney, T. E., Braviak, L., Tato, C. M. & Khatri, P. Genome-wide expression for diagnosis of pulmonary tuberculosis: a multicohort analysis. Lancet Respir. Med. 4, 213–224 (2016).

33. Andres-Terre, M. et al. Integrated, Multi-cohort Analysis Identifies Conserved Transcriptional Signatures across Multiple Respiratory Viruses. Immunity 43, 1199–1211 (2015).

34. Ennis, F. A. & Meager, A. Immune interferon produced to high levels by antigenic stimulation of human lymphocytes with influenza virus. J. Exp. Med. 154, 1279–1289 (1981).

35. Baserga, R. THE RELATIONSHIP OF THE CELL CYCLE TO TUMOR GROWTH AND CONTROL OF CELL DIVISION: A REVIEW. Cancer Res. 25, 581–595 (1965).

36. Diehl, A. D., Lee, J. A., Scheuermann, R. H. & Blake, J. A. Ontology development for biological systems: immunology. Bioinformatics 23, 913–915 (2007).

37. Manning, C. D. & Schütze, H. Foundations of statistical natural language processing. (MIT Press, 1999).

38. Ballouz, S., Pavlidis, P. & Gillis, J. Using predictive specificity to determine when gene set analysis is biologically meaningful. Nucleic Acids Res. gkw957 (2016). doi:10.1093/nar/gkw957

39. Blake, J. A. Ten quick tips for using the gene ontology. PLoS Comput. Biol. 9, e1003343 (2013).

40. Hartung, M., s, A. G. & Rahm, E. Rule-based Generation of Diff Evolution Mappings between Ontology Versions. CoRR abs/1010.0122, (2010).

41. Resnik, P. Using Information Content to Evaluate Semantic Similarity in a Taxonomy. in In Proceedings of the 14th International Joint Conference on Artificial Intelligence 448–453 (1995).

42. Harispe, S., Ranwez, S., Janaqi, S. & Montmain, J. The semantic measures library and toolkit: fast computation of semantic similarity and relatedness using biomedical ontologies. Bioinforma. Oxf. Engl. 30, 740–742 (2014).

